# Isolation and Divergence of *Peromyscus melanotis* Populations Across the Madrean Sky Islands in Arizona

**DOI:** 10.1101/2022.04.11.487881

**Authors:** Brendan B. Larsen, Hans W. Otto, Sophie Gryseels, Michael Worobey

## Abstract

The Madrean sky islands have been studied for decades due to their high biodiversity, which results from the intersection of biomes and their role as refugia for populations isolated on mountain tops during the last ice age. There has been controversy and confusion about the identification of *Peromyscus sp*. found in the montane forests on these sky islands, which is often assumed to be the widespread and ubiquitous *P. maniculatus*. Here, we provide mitochondrial phylogenetic data suggesting that all individuals of *Peromyscus* captured on three isolated mountains in southern Arizona are *Peromyscus melanotis*, a species previously thought endemic to Mexico. Furthermore, with molecular clock analyses on two mitochondrial loci we show these populations have been isolated from each other for ∼11,000-50,000 years, corresponding to the transition from the last ice age. These isolated populations represent important conservation targets due to habitat loss. In addition, we suggest that future genomic and ecological research is warranted to better understand these unique populations.

## Introduction

The Madrean sky islands are one of the most biodiverse regions in North America, characterized by a series of isolated mountain ranges spanning from Arizona and New Mexico, south into Mexico where they merge with the Sierra Madre Occidental (DeBano 1999). These mountain ranges are often surrounded by low elevation desert that act as a filter route for many species of flora and fauna adapted to high elevation montane forest. They have been termed ‘sky islands’ because they are disjunct high elevation montane habitats surrounded by desert “seas”. Evidence from ancient packrat middens spanning the last ∼40,000 years suggest that ∼26,000 years ago, during the last glacial maximum (LGM), these sky islands were entirely connected by woodlands (Betancourt et al. 1990).

*Peromyscus*, in particular *Peromyscus maniculatus*, are a model species for a wide range of biological disciplines and are one of the most ubiquitous rodent species in North America (Bedford and Hoekstra 2015). In Arizona they can be found in nearly all habitats, ranging from dry hot desert to coniferous forests at high elevation (Hoffmeister 1986). In recent years the taxonomy and evolutionary relationship of *Peromyscus* has been refined as genetic work has overturned putative relationships based solely on morphology (Dragoo et al. 2006; Bradley et al. 2007; Platt et al. 2015).

*Peromyscus melanotis*, or the Black-eared deer mouse, was described in Osgood’s original treatment of *Peromyscus* species as being endemic to high elevation (>2000m) areas of the Sierra Madre Occidental in Mexico (Osgood 1909). In the northern parts of their geographical distribution, *P. melanotis* may be confused with *P. maniculatus* based on morphological characteristics in size and coloration. There is considerable overlap in external measures, weight, dental formula (identical), and cranial measurements between these species and often cannot be used as distinguishing characteristics. Interestingly, karyotyping of montane *Peromyscus* populations from three Madrean sky islands including the Santa Catalina Mountains, Pinaleño Mountains, and Chiricahua Mountains (hereafter, study sites) in southern Arizona revealed they share features with *P. melanotis* and not lowland *P. maniculatus* (Bowers et al. 1973). Prior to this study, the closest record of *P. melanotis* to Arizona was ∼177 km in Chihuahua, Mexico (Hoffmeister 1986). Additional research with captive breeding colonies provided evidence that montane *Peromyscus* from these Madrean sky islands were unable to produce viable offspring with lowland *P. maniculatus* (Bowers et al. 1973; Bowers 1974). In a later analysis, Hoffmeister discusses the possibility of *P. melanotis* being present in Arizona but rejected this possibility due to absence of any definitive morphological character differences between the montane *Peromyscus sp*. populations with *P. maniculatus* (Hoffmeister 1986). It is possible that the large influence Hoffmeister had on mammalogy research in Arizona led to little further exploration of this issue.

Confusion about the status of *P. melanotis* in Arizona persists today. In the Mammalian Species account (Álvarez-Castañeda 2005), *P. melanotis* is listed as a native species of Arizona. However, authors cite a review from 1989 that fails to mention Hoffmeister’s conclusion based on morphology, and instead use Bowers karyotyping and breeding research as evidence for species presence. Additional confusion results from mammal lists that include *P. melanotis* as a native mammal of the United States, and others omit the species entirely (Bowers et al. 2007; Ammerman et al. 2014). Furthermore, DNA collected from *Peromyscus* on the upper peaks of the Chiricahua mountains is referenced as *P. melanotis* (Gering et al. 2009). These data confirm that the DNA is distinct from *P. maniculatus* genotypes collected in Colorado (Gering et al. 2009), and that mitochondrial sequences from rodent individuals from the Chiricahua mountains cluster with *P. melanotis* and not *P. maniculatus*, but the significance of this finding was not discussed.

Given the lack of definitive morphological differences between *P. melanotis* and *P. maniculatus*, it is understandable there is confusion about the status of these species in southern Arizona. Here, we attempt to resolve some of these long-standing questions by collecting and analyzing mitochondrial sequence data of *Peromyscus* rodents from the three study sites for which Bowers work originally suggested as belonging to *P. melanotis*. Based on the geographic isolation of these three montane habitats, we hypothesized that these same mitochondrial sequences would show levels of divergence consistent with isolation since the last glacial maximum.

## Methods

Our study was conducted in the Santa Catalina, Pinaleño, and Chiricahua Mountains ranges in the Coronado National Forest in Pima, Pinal, Cochise, and Graham Counties Arizona (GPS coordinates and elevation of all trapping locations are given in Supplementary Table 1). We trapped a range of elevations from 2334-2725m. Within Coronado National Forest there are several biotic communities characterized by unique flora and fauna. Our study sites took place in a transition zone known as Petran montane conifer forest. Common vegetation in this zone includes ponderosa pine (*Pinus ponderosa*), southwestern white pine (*Pinus strobiformis*), Douglas fir (*Pseudotsuga menziesii*), Gambel oak (*Quercus gambelii*), and New Mexican locust (*Robinia neomexicana*). Soil was friable, and abundant leaf litter was present throughout each study site.

To capture *Peromyscus*, we set ∼100 Sherman live traps (7.6 cm x 7.6 cm x 25.4 cm) for two consecutive nights from 12 May to 30 June 2019 at each study site (2 per mountain). We strategically placed traps near hollow logs in coniferous forest, baited traps with rolled oats and mixed birdseed, set them near dusk, and checked them early the following morning. For mammals captured, we recorded species, sex, age (juvenile or adult), reproductive condition (males-scrotal or nonscrotal; females-pregnant, lactating, postlactating, or nonreproductive), and weighed individuals with a Pesola spring scale (Rebmattli 19, CH-6340, Baar, Switerland). From May to June, we recorded 21 captures of *Peromyscus* that based on morphology could not be identified as either *P. melanotis* or *P. maniculatus*. Other species captured at study sites included *Reithrodontomys megalotis, Sigmodon arizonae, Neotoma mexicana*, and *Peromyscus boylii*. For all individuals identifiable as either *P. melanotis* or *P. maniculatus*, a clip of ear tissue was collected using sterile surgical scissors. Ear clips aided in reidentification of individuals, and were stored in RNAlater (ThermoFisher, Waltham, MA, USA) and then placed in a −80°C freezer within 48 hours of collection. We released all individuals at points of capture. All capture and animal handling procedures were approved by the University of Arizona Institutional Animal Care and Use Committee (IACUC 15-583), and the Arizona Department of Game and Fish (permit # SP506475).

To extract DNA, ear clips were removed from the RNAlater and washed briefly with 1x PBS (ThermoFisher, Waltham, MA, USA). Ear clip tissue was then processed with the E.Z.N.A Tissue DNA kit (Omega Bio-tek, Norcross, GA, USA). We amplified the complete mitochondrial cytochrome-*b* (cyt*b*) coding region based on previously described primer sets (Naidu et al. 2012). 1 ng of DNA was used as a template. 25 µL PCR were set up with 1X PCR Buffer (NEB, Ipswich, MA, USA) containing 1.5 mM MgCl_2_, with an additional 1 mM MgCl_2_. We also added 0.5 µM of forward and reverse primer, 200 µM dNTP, and 0.625 Taq (NEB, Ipswich, MA, USA). PCR cycling conditions were an initial denaturation at 95°C for 1 minute, followed by 35 cycles at 95°C for 30 s, annealing at 55°C for 30 s, and extension at 68°C for 1 min 30s. DNA from the mitochondrial control region D-loop region was amplified using the primers D_F (5’-CATCAACACCCAAAGCTGATATTC-3’) and D_R (5’-CAAAAGGTTTGGTCCTGGCCTTAT-3’). PCR conditions were identical as above, except the annealing temperature was changed to 52°C. All PCR included negative controls, and PCR products were visualized on a 1.5% Agarose gel. All amplicons were prepared for sequencing with the DNA Clean & Concentrator Kit (Zymo Research, Irvine, CA, USA) and were sequenced by Sanger in both directions on an Applied Biosystems 3730 at the University of Arizona Genetics Core. Sanger chromatograms were assembled in Geneious v8.1.5 (Biomatters, Auckland, New Zealand).

Sequence alignments were made with MAFFTv7.475 (Katoh and Standley 2013) and manually trimmed to remove primer sequence. Phylogenies were inferred with MrBayes v3.2.7a (Ronquist and Huelsenbeck 2003) using a GTR substitution model and Gamma rate variation for 10,000,000 generations. Trees were sampled every 1000 generations and the first 20% were discarded as the burn-in.

We performed molecular clock analyses on individual cyt*b* and control region alignments. We used a GTR substitution model with 4 rate categories, and a strict clock. We used three different rodent mitochondria substitution rate priors to estimate the divergence of these sky island populations. First, for the cyt*b* clock rate, we used a prior with a normal distribution with a mean of either 6e-8 or 3.2e-7 subs/site/year and a standard deviation of 7e-9 or 4e-8, respectively (Herman and Searle 2011; Martínková et al. 2013; Platt et al. 2015). For the control region clock rate we used a prior with a normal distribution with a mean of 5.6e-8 subs/site/year and a standard deviation of 7e-9 (Goios et al. 2007).

## Results

### Identity of rodents captured

We trapped at two separate locations on each of the mountain ranges that make up this study. A topographic map of S Arizona and NW Mexico with trapping locations is shown in Figure 1. Because several species of *Peromyscus* (*e*.*g*., *P. boylii*) are sympatric at high elevations within this region, we only extracted ear clippings of *Peromyscus* species that were identifiable as either *P. melanotis* or *P. maniculatus*. From these, we extracted DNA and amplified cyt*b* sequences from 12 individuals captured in the Santa Catalina Mountains, five individuals captured in the Chiricahua Mountains, and four individuals captured in the Pinaleño Mountains. All clustered with *P. melanotis* mitochondrial sequences and none with *P. maniculatis*. We then compared these cyt*b* sequences with all available *P. melanotis* cyt*b* sequences in GenBank (including a previously-sequenced individual from the Chiricahuas, EU574690, Supplementary Table 2, (Gering et al. 2009)). A Bayesian phylogeny based on this set of *P. melanotis* cyt*b* sequences and other *Peromyscus* reference outgroup sequences shows the three Arizona sky island populations each form distinct clades within the *P. melanotis* diversity (Figure 2).

**Figure 1.**
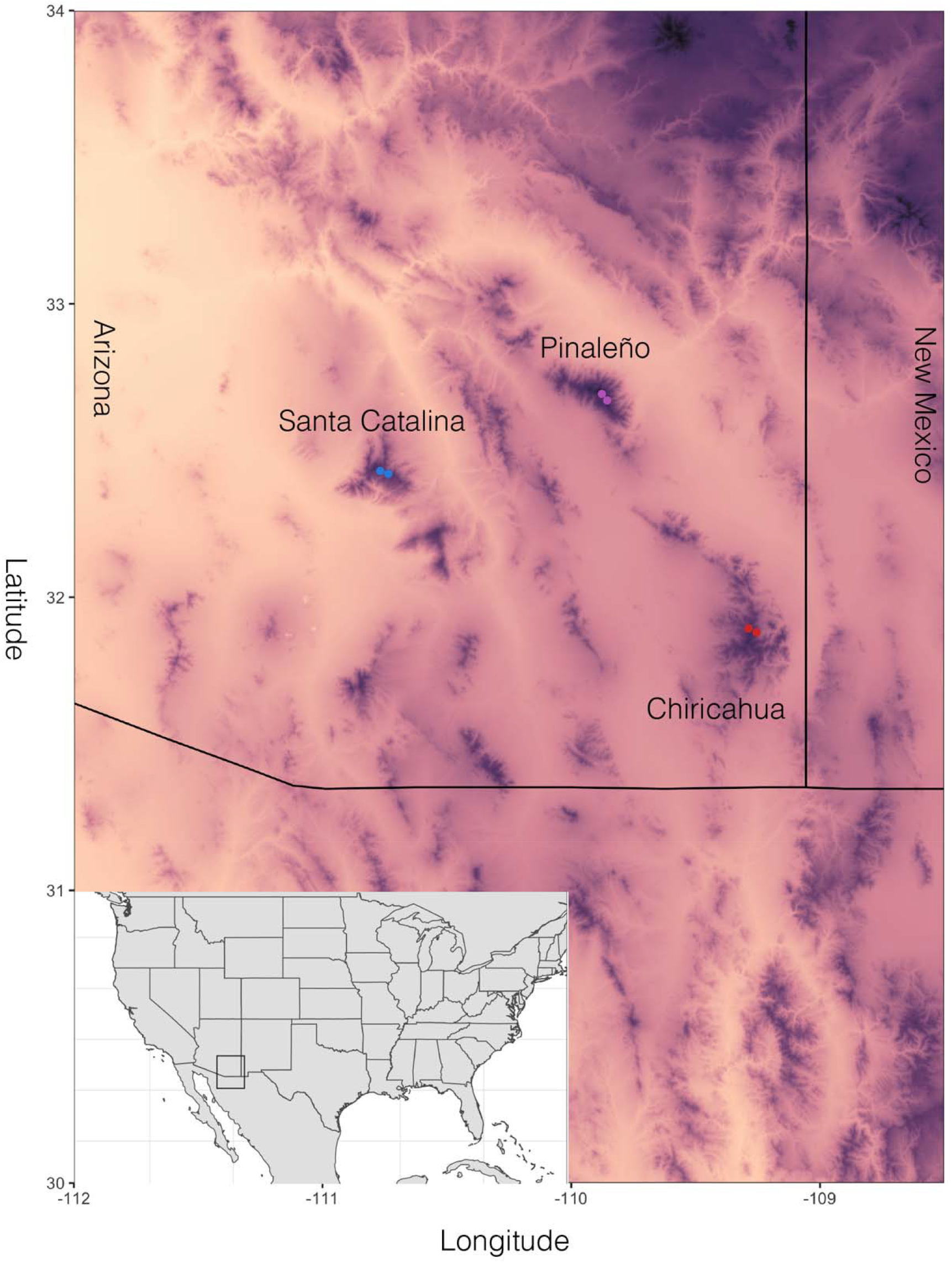
Topographic map of southeastern Arizona and trapping sites. Darker colors represent higher elevations, while lighter colors represent lower elevations. Colored dots represent trapping locations used in this study.

**Figure 2.**
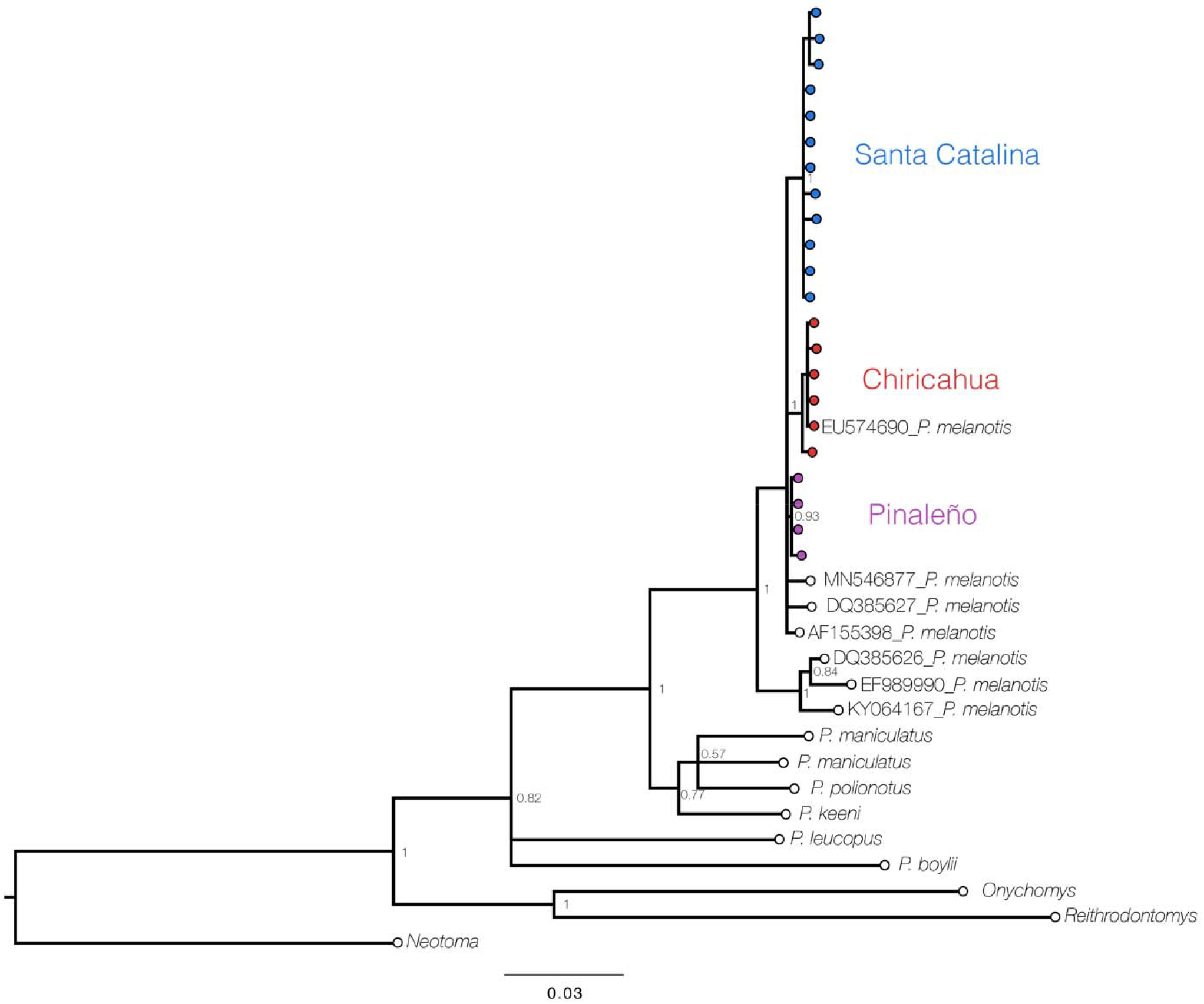
Bayesian phylogeny based on cyt*b* of representative *Peromyscus* species and their allies. The Arizona sequences are colored by the mountain range they come from. Genbank accession numbers for the P. melanotis sequences are included in the name for reference. Numbers at nodes are posterior probabilities.

### Phylogenetic relationships among the three sky island populations

The phylogeny based on only the cyt*b* sequence information resulted in a polytomy between the three sky island populations. In order to better resolve the relationship between these three populations we concatenated the control region sequences with the cyt*b* sequences to increase the amount of sequence data. This analysis was restricted to our sky island samples because only cyt*b* sequence data is available in GenBank from previously sampled *P. melanotis* from Mexico and the Chiricahuas. Bayesian phylogenetic analysis revealed the Chiricahua and Pinaleño mountain populations are sister to each other (Figure 3, Posterior Probability=0.7), with the Santa Catalina mountain population being the outgroup.

**Figure 3.**
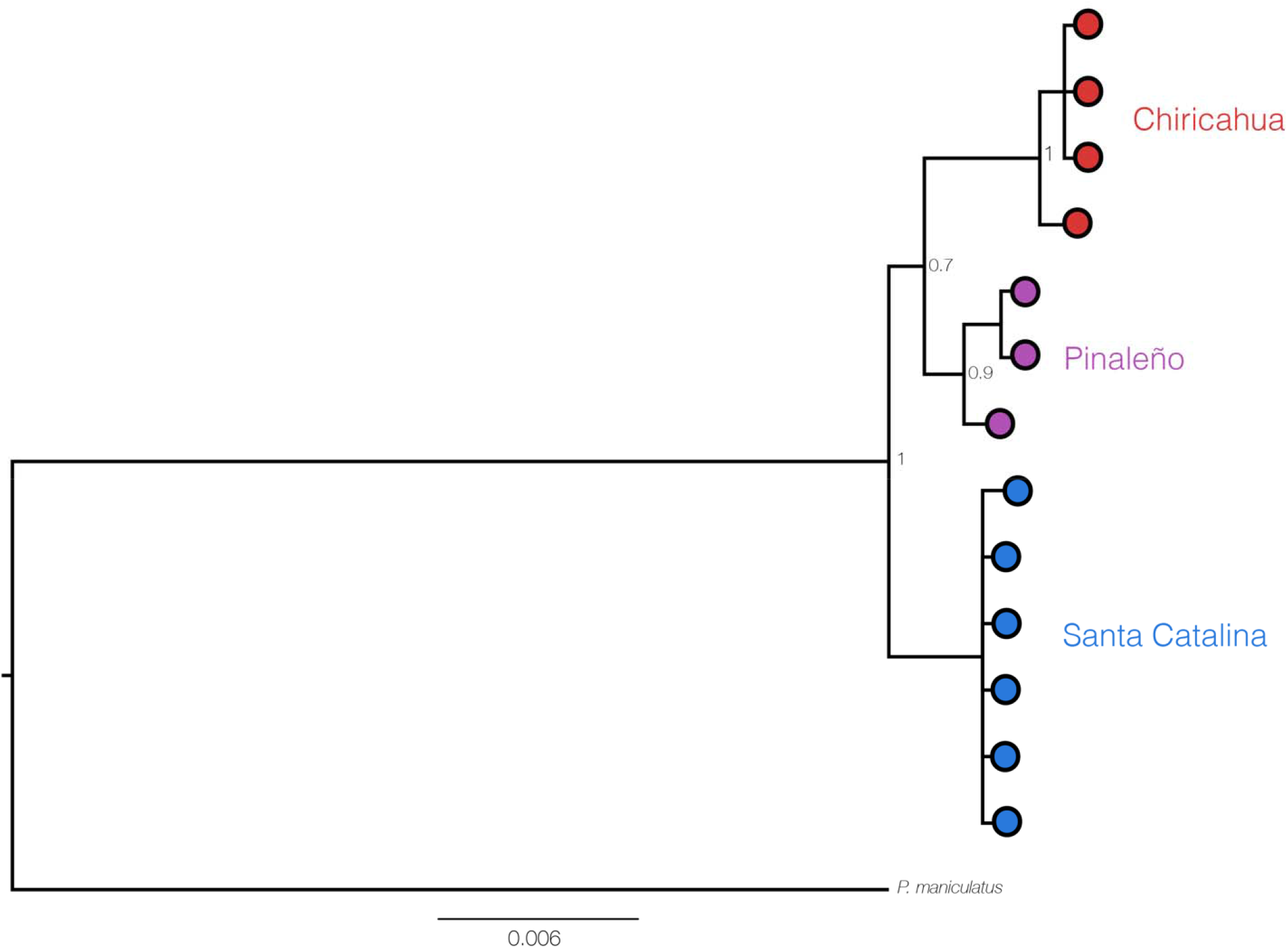
Bayesian phylogeny of the concatenated cyt*b /*control region sequences for the three Arizona sky island populations and an outgroup reference sequence of P. maniculatus downloaded from Genbank (NC_039921.1). Numbers at nodes are posterior probabilities.

### Divergence estimates for three sky island populations of Peromyscus

To estimate the divergence time between these three populations, we performed molecular clock analyses. Because estimates are highly sensitive to priors and the loci used, we sequenced an additional stretch of mitochondrial DNA, the control region, for which there is additional data on the substitution rate derived from the literature. We then used two different estimates of the rodent cyt*b* substitution rate, and the substitution rate estimate of the control region. First, we only used the Arizona cyt*b* sequences and used a substitution rate with a normal distribution of 6e-8 subs/site/year (stdev 7e-9) based on previous estimates (Platt et al. 2015). The median estimated tMRCA for the three sky island populations was 59,997 years before present (ybp) (95% HPD 28,651-102,190). Next, we used a faster cyt*b* substitution rate based on ancient DNA studies of *Microtus* with a prior substitution rate normal distribution of 3.2e-7 subs/site/year (stdev 4e-8) (Herman and Searle 2011; Martínková et al. 2013). The median estimated tMRCA was 11,218 ybp (95% HPD 5472 - 19576). Finally, using the mitochondrial control region and independent substitution rate prior normal distribution of 5.6e-8 subs/site/year (stdev 7e-9) (Goios et al. 2007), the median tMRCA of the Arizona sky island populations was 52,174 ybp (95% HPD 20,909-97,255). These estimates are summarized in Figure 4.

**Figure 4.**
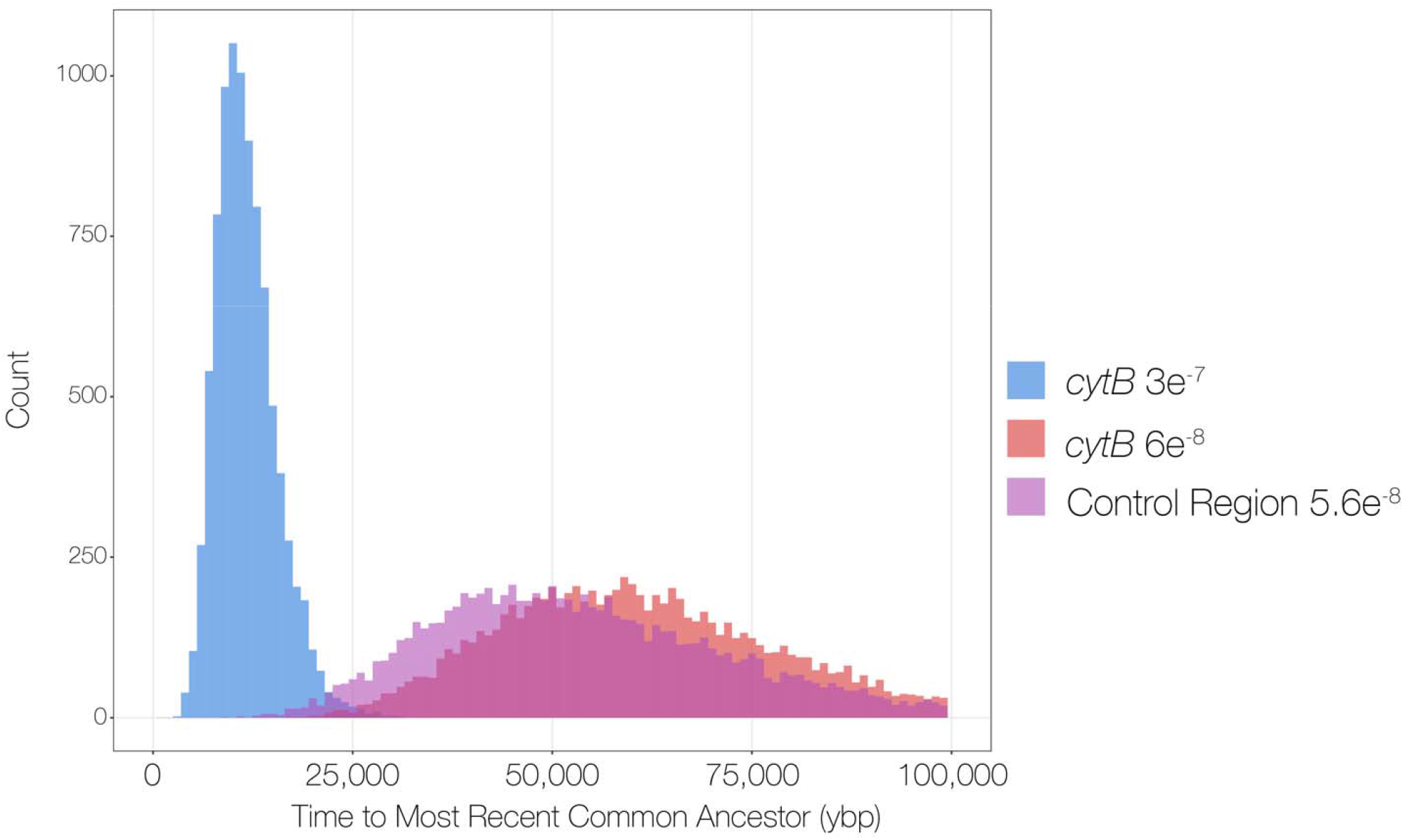
Bayesian posterior distributions for the time to the most recent common ancestor of the three Arizona sky island populations colored by locus and mean substitution rate prior used.

## Discussion

Through genetic barcoding we resolve prior confusion and establish that *P. melanotis* is present on three sky islands in southern Arizona, as suggested by Bowers et al. 1974. We also show these three populations have been isolated for ∼11,000-55,000 years, coinciding with the last glacial maximum, with evidence that the Pinaleño and Chiricahua mountain populations diverged more recently than the Santa Catalina mountain population. As the climate warmed, these populations became isolated and have been evolving independently since the last glacial maximum.

One of the major limitations of our study was the use of only mitochondrial data for species identification. It is possible that regions of the mitochondria are introgressed from *P. melanotis* while the nuclear genome is derived from *P. maniculatus*. Introgression of mitochondrial haplotypes has been observed for other mammal lineages, such as bears (Miller et al. 2012), and even other species of *Peromyscus* (Garcia-Elfring et al. 2017). Further sequencing of the nuclear genome should be performed on these Arizona sky island populations to confirm our results.

Although our timing estimates are based on limited mitochondrial sequence data, we argue the time scale is realistic. We used previously published cyt*b* and control region substitution rates to calibrate our molecular clock analysis. It is important to note there is no single established rodent mitochondrial substitution rate. These rates can vary greatly by species and especially depending on the time scale used (Ho et al. 2005; Henn et al. 2009). In order to address this issue, we selected three different rodent substitution rates that are based on different lines of evidence to come to a range of possibilities. The first estimate of 6e-8 subs/site/year for cyt*b* is based on deeper time scales and rodent fossil calibrations, and likely represents the maximum time estimate (Platt et al. 2015). The second estimate of 3.2e-7 subs/site/year for cyt*b* is based on a much shallower time estimate from the rodent genus *Microtus* based on ancient DNA (Herman and Searle 2011; Martínková et al. 2013). Finally, the third estimate of 5.6e-8 subs/site/year for the control region is based on ∼100 years divergence of *Mus musculus* domesticated rodent stocks. All three of the substitution rate estimates result in divergence times that overlap with the most recent glaciation cycle. These substitution rate estimates, gathered from different rodent species and estimated using different methods and time scales, converge on these populations diverging within the last 100,000 years, and likely correspond to the last glacial maximum 26,000 years ago.

Although we were able to demonstrate this species was present on three sky islands, we were not able to address the overall geographic extent of this species in the United States. It is likely that it also occurs in high-elevation forests on the other, non-sampled Madrean sky islands of southern Arizona and New Mexico. The high elevation pine forests of the Santa Rita and Huachuca mountains are similar habitats as those sampled in this study. One outstanding question is if this species makes it farther north into areas of the Mogollon Rim or other sky islands in Arizona and New Mexico, or if the northern limit of this species stops at the Pinaleño and Santa Catalina mountains. Due to morphological similarities with *P. maniculatus* it is essential to sequence a DNA barcode to identify the species. It is worth noting that some *Peromyscus* have been DNA barcoded from the Mogollon Rim and are confirmed as *P. maniculatus*. This does not rule out the possible occurrence of *P. melanotis* in these regions. However, since many Sierran Madrean neotropical species reach their northern limit in the Madrean sky islands (Warshall 1994) it is likely we detected the northern limit of *P. melanotis*.

These isolated populations in the Madrean sky islands in Arizona likely represent the total population of *P. melanotis* in the United States. Therefore, they are important targets for future conservation efforts. Human activities including overgrazing by livestock, mining encroachment, and fire suppression directly threaten biotic communities in this area. The preeminent threat for these species is the increase in catastrophic wildfires due to climate change and human interventions (Coe et al. 2012). In 2011, the Horseshoe II fire burned 220k acres in Chiricahua mountains, the Nuttal fire in 2004 and the Frye Fire in 2017 burned much of the upper reaches of the Pinaleño mountains, and in 2020 the Bighorn fire burned ∼120k acres of the Santa Catalina mountains. Nothing is known about how *P. melanotis* recovers and colonizes burned habitats following wildfires. Population surveys of *P. melanotis* in the sky islands would be informative to determine whether this species should be considered for threatened or endangered status in the United States.

With these data we add to the taxonomic list of Arizona Madrean sky island biota that have been evolutionary isolated during recent glacial cycles. Surprisingly, several studies show a deeper divergence pattern (mya) in other taxa compared to our analyses (Masta 2000; Tennessen and Zamudio 2008; Wiens et al. 2019), though for Mexican Jays similar recent divergences were inferred (McCormack et al. 2008). More extensive genome sequencing of *P. melanotis* and the inclusion of sky island populations in Mexico would help explain the complex demographic and migration patterns that have occurred over the past 100,000 years for this species. *P. melanotis* represents a good candidate for future genome analyses in Arizona to study parallel evolution. The populations we sampled have been evolving independently in nearly identical habitats for the past 11k-55k years, and are closely related to *P. maniculatus*, for which many genomic resources are available.

One final interesting observation from this study is that although our trapping numbers were limited (*n=*21), we never caught and genotyped *P. maniculatus* at the high elevations of the three sky islands we studied. *P. maniculatus* is known to occupy high elevations (∼4300 m in Colorado) in portions of its range, and the coniferous forest habitat in our study sites should be ideal for this species (Hoffmeister 1986; Gering et al. 2009). Possibly *P. maniculatus* is present but *P. melanotis* is dominant. We did not trap or genotype extensively enough to determine the lower elevational limit of *P. melanotis* in Arizona, or to what extent *P. melanotis* and *P. maniculatus* are sympatric. Future genomic and ecological research is warranted to better understand the distributions and habitat preferences of these unique populations.

## Acknowledgements

We thank the Arizona Department of Game and Fish for allowing us permission to conduct research in Coronado National Forest. We also thank K. Chenard, D. Davison and A. Potticary for assistance with fieldwork. B.B.L. was supported by funding from the NSF GRFP, Society for the Study of Evolution Rosemary Award, and the University of Arizona Galileo Scholar program. Funding was provided to M.W. from the David and Lucile Packard Foundation. SG was supported by an OUTGOING [PEGASUS]^2^ Marie Skłodowska-Curie Fellowship (12T1117N) of the Research Foundation – Flanders (FWO).

## Supplementary data

**Supplementary Table 1**. GPS coordinates for locations that were trapped

**Supplementary Table 2**. Genbank accession IDs used in analysis

---

**Supplementary Table 1.**
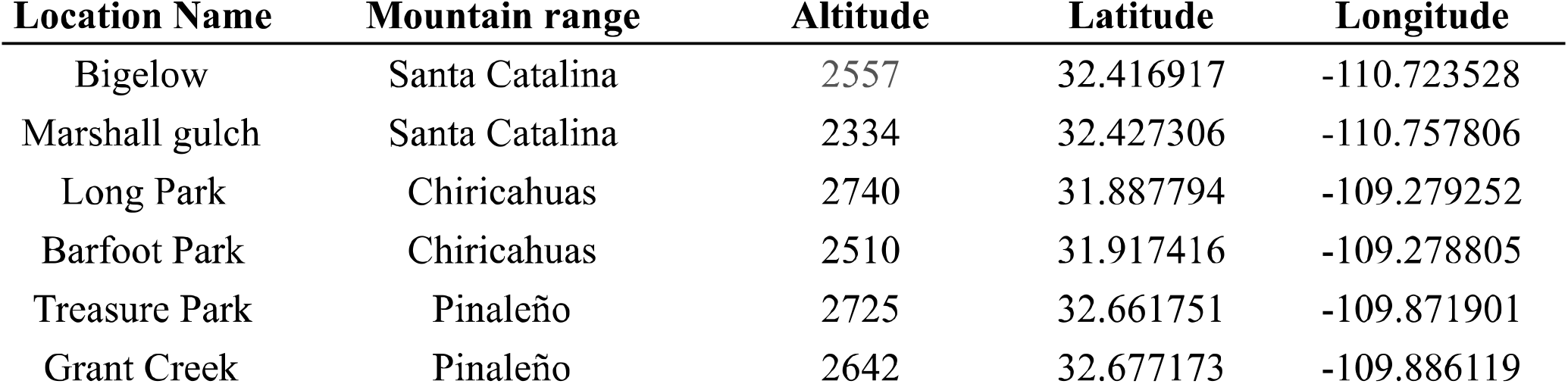
Locations of all trapping locations.

| Location Name | Mountain range | Altitude | Latitude | Longitude |
| --- | --- | --- | --- | --- |
| Bigelow | Santa Catalina | 2557 | 32.416917 | -110.723528 |
| Marshall gulch | Santa Catalina | 2334 | 32.427306 | -110.757806 |
| Long Park | Chiricahuas | 2740 | 31.887794 | -109.279252 |
| Barfoot Park | Chiricahuas | 2510 | 31.917416 | -109.278805 |
| Treasure Park | Pinaleno | 2725 | 32.661751 | -109.871901 |
| Grant Creek | Pinaleno | 2642 | 32.677173 | -109.886119 |

**Supplementary Table 2.**
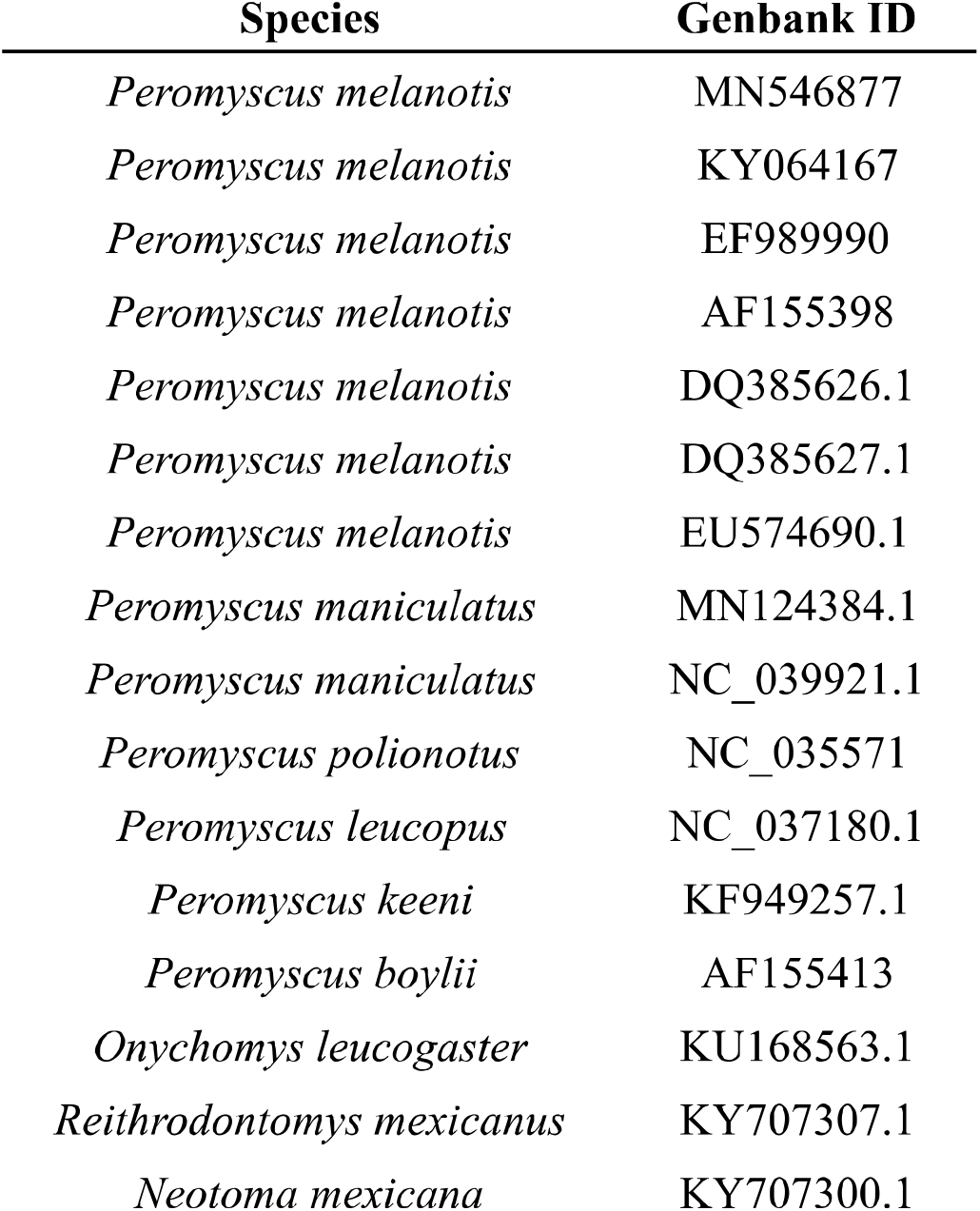
Genbank accessions used in paper.

