## Supplemental Data 1 for "Isolation and Divergence of *Peromyscus melanotis* Populations Across the Madrean Sky Islands in Arizona"

| Location Name | Mountain range | Altitude | Latitude | Longitude |
|---|---|---|---|---|
| Bigelow | Santa Catalina | 2557 | 32.416917 | -110.723528 |
| Marshall gulch | Santa Catalina | 2334 | 32.427306 | -110.757806 |
| Long Park | Chiricahuas | 2740 | 31.887794 | -109.279252 |
| Barfoot Park | Chiricahuas | 2510 | 31.917416 | -109.278805 |
| Treasure Park | Pinaleño | 2725 | 32.661751 | -109.871901 |
| Grant Creek | Pinaleño | 2642 | 32.677173 | -109.886119 |
