## Supplemental Data 2 for "Isolation and Divergence of *Peromyscus melanotis* Populations Across the Madrean Sky Islands in Arizona"

| Species | Genbank ID |
|---|---|
| *Peromyscus melanotis* | MN546877 |
| *Peromyscus melanotis* | KY064167 |
| *Peromyscus melanotis* | EF989990 |
| *Peromyscus melanotis* | AF155398 |
| *Peromyscus melanotis* | DQ385626.1 |
| *Peromyscus melanotis* | DQ385627.1 |
| *Peromyscus melanotis* | EU574690.1 |
| *Peromyscus maniculatus* | MN124384.1 |
| *Peromyscus maniculatus* | NC_039921.1 |
| *Peromyscus polionotus* | NC_035571 |
| *Peromyscus leucopus* | NC_037180.1 |
| *Peromyscus keeni* | KF949257.1 |
| *Peromyscus boylii* | AF155413 |
| *Onychomys leucogaster* | KU168563.1 |
| *Reithrodontomys mexicanus* | KY707307.1 |
| *Neotoma mexicana* | KY707300.1 |
